# Huntington’s disease-associated ankyrin repeat palmitoyl transferases are rate-limiting factors in lysosome formation and fusion

**DOI:** 10.1101/2025.02.06.636816

**Authors:** Győző Szenci, Attila Boda, Anikó Nagy, Zsombor Szőke, Gergely Falcsik, Tibor Kovács, Péter Lőrincz, Gábor Juhász, Szabolcs Takáts

## Abstract

Protein palmitoylation in the Golgi apparatus is critical for the appropriate sorting of various proteins belonging to secretory and lysosomal systems, and defective palmitoylation can lead to the onset of severe pathologies. HIP14 and HIP14L ankyrin repeat-containing palmitoyl transferases were linked to the pathogenesis of Huntington’s disease, however, how perturbation of these Golgi resident enzymes contributes to neurological disorders is yet to be understood. In this study, we investigated the function of Hip14 and Patsas - the Drosophila orthologs of HIP14 and HIP14L respectively – to uncover their role in secretory and lysosomal membrane trafficking. Using larval salivary gland, a well-established model of the regulated secretory pathway, we found that these PAT enzymes equally contribute to the proper maturation and crinophagic degradation of glue secretory granules by mediating their fusion with the endo-lysosomal compartment. We also revealed that Patsas and Hip14 are both required for lysosomal acidification and biosynthetic transport of various lysosomal hydrolases, and we demonstrated that the rate of secretory granule-lysosome fusion and subsequent acidification positively correlates with the level of Hip14. Furthermore, Hip14 is also essential for proper lysosome formation and neuronal function in adult brains. Finally, we found that the over-activation of lysosomal biosynthetic transport and lysosomal fusions by the expression of the constitutively active form of Rab2 could compensate for the lysosomal dysfunction caused by the loss of Patsas or Hip14 both in larval salivary glands and neurons. Therefore, we demonstrated that ankyrin repeat palmitoyl transferases may act as rate-limiting factors in lysosomal fusions and provide genetic evidence that defective protein palmitoylation and the subsequent lysosomal dysfunction can contribute to the onset of Huntington’s disease-like symptoms.

**Author Summary:** Growing body of evidence suggests that decreased activity of HIP14 and HIP14L palmitoyl transferases caused by accumulation of mutant Huntingtin and the subsequent alterations in protein palmitoylation play a critical role in the onset of Huntington’s disease (HD). However, which cellular processes are perturbed and eventually lead to the emergence of HD due to impaired palmitoylation is still poorly understood. In our study, we used a *Drosophila* model to uncover the role of Hip14 and Patsas (the fly ortholog of HIP14L) in secretory and lysosomal membrane trafficking. We found that silencing of these transferases in larval salivary glands equally disrupts secretory granule lysosome-fusion, their subsequent acidification and also the trafficking of lysosomal hydrolases. While overexpression of Hip14 resulted in the acceleration of these processes. We also observed that neuron-specific loss of Hip14 not only perturbs lysosome formation but also results in a progressive decline in neuromuscular functions. Importantly, both lysosomal and neuronal defects emerging in Hip14 and Patsas deficient backgrounds could be restored by hyperactivation of Rab2 GTPase mediated lysosome formation and fusion. These findings suggest that Hip14 and Patsas protect from the onset of HD like symptoms by acting as rate-limiting factors of lysosome formation and fusion.

## Introduction

Palmitoylation is a reversible post-translational lipid modification that alters the substrate proteins’ hydrophobicity thus affecting their intracellular trafficking, membrane association, stability and interactions with other proteins. Regulators of intracellular signaling and transport are over-represented among these substrate proteins [1, 2], hence altered palmitoylation is implicated in the emergence of various diseases such as cancer, diabetes and neurological disorders [3]. The transfer of the palmitate chain to Cys residues of substrate proteins is catalyzed by DHHC (Asp-His-His-Cys) palmitoyl acyltransferase (PAT) enzymes, consisting of 4-6 transmembrane domains and a cytosolic Cys-rich domain (CRD) encompassing the eponymous DHHC catalytic motif. In addition, mammalian DHHC17/HIP14 (Huntingtin interacting protein 14) and DHHC13/HIP14L (Huntingtin interacting protein 14 like) harbor an additional N-terminal Ankyrin repeat domain (ANK), that interacts with and regulates the subcellular localization of several substrates involved in neuronal function, including the Huntington disease-associated protein Huntingtin (HTT) [4–6]. In Huntington’s disease (HD), the CAG nucleotide expansion in HTT gene causes a poly-Q expansion in HTT protein which decreases its interaction with HIP14 and HIP14L, consequently, their enzymatic activity is lowered [5, 7, 8], presumably resulting in the mislocalization of multiple synaptic proteins [9]. In line with this, mice lacking HIP14 or HIP14L develop neuropathological phenotypes akin to HD [8, 10, 11]. The delivery of synaptic proteins is linked to the secretory pathway. Sorting of the secretory proteins with different cellular destinations (plasma membrane, lysosome, secretory granules) takes place at the trans-Golgi network [12] and palmitoylation has been implicated as a key regulator of anterograde, plasma membrane-directed transport of proteins [13]. Since both HIP14 and HIP14L are Golgi resident enzymes, the mislocalization of synaptic proteins due to defective palmitoylation suggests that these PATs may have a critical role in maintaining proper protein sorting in the Golgi. However, how these PATs regulate post-Golgi membrane trafficking pathways is largely unexplored.

*Drosophila* is an excellent and popular in vivo experimental system for modeling human diseases (like HD or other neurodegenerative disorders) and understanding the underlying alterations in genetics and cell biology. Moreover, the salivary gland of late L3 staged *Drosophila* larvae also became a powerful platform for studying the genetic regulation of post-Golgi trafficking, and particularly the complicated relationship of secretory and lysosomal pathways. Larval salivary gland cells synthesize and secrete high amounts of mucous Sgs (salivary gland secretion) or glue proteins in response to the molting hormone ecdysone [14–16]. The glue proteins at the trans-Golgi network are packaged into immature secretory granules [17, 18], which undergo a complex maturation process involving a significant increase in size by homotypic fusions [19–22], progressive acidification [23–26], the profound remodeling of the secretory material [23–26] and the acquisition of membrane proteins required for exocytosis [21, 27]. This is ensured by a series of fusion events between the maturing secretory granules and lysosomes that control the quantity and quality of the secretory material along the entire secretory pathway [28]. Hence, fusion of secretory granules with a heterogenous non-degradative lysosomal population is critical for their maturation [24–26], while, the excess or abnormal secretory granules fuse with degradative lysosomes to be selectively degraded by crinophagy [25, 26, 29, 30].

The *Drosophila* genome encodes two ANK domain-containing PATs: Hip14 (CG6017) and Patsas (CG6618), which are abundant in nervous tissues and localized to the Golgi apparatus [31]. Consistent with the mammalian results, Hip14 is essential for the proper development and efficient synaptogenesis and synaptic transmission by regulating the delivery of cysteine string protein (CSP) and synaptosome-associated protein 25 (Snap-25) by direct palmitoylation [32–34]. In contrast, Patsas is still poorly characterized. This raises the possibility that Hip14 (and potentially Patsas) has a conserved function in synaptic trafficking, but how these Golgi localized ANK PATs control post-Golgi secretory pathways at cellular and molecular levels is still elusive. Here we use the *Drosophila* larval salivary gland to understand the role of Hip14 and Patsas in the regulated secretory pathway and uncover their role in secretory granule-lysosome fusions and highlight its potential implications on neuronal function.

## Results

### 1. Patsas and Hip14 are essential for proper crinophagic degradation of the content of secretory granules and their fusion with PI3P-positive endosomes

We chose the salivary gland of *Drosophila* larvae as a model system to study the role of ANK PATs in post-Golgi trafficking. Since maturation or crinophagic degradation of secretory granules requires the fine-tuned interplay of the secretory and lysosomal systems, these serve as good indicators of the integrity of post-Golgi trafficking. First, we examined the potential role of the Patsas and Hip14 PAT enzymes in crinophagic degradation. At the prepupal stage (pp), most glue granules are already secreted, and the non-secreted granules trapped in the cell fuse with degradative lysosomes to form crinosomes. To assess the efficiency of crinophagic degradation we carried out a crinophagic flux assay by expressing the N-terminally GFP- and dsRed-tagged Sgs3 glue protein in secretory gland cells. The two reporters are equally sorted into the forming glue granules, so these appear as GFP and dsRed double positive structures. After fusing with acidic lysosomes the secretory granules are transformed into degradative crinosomes in which the highly acidic milieu leads to the quenching of GFP (but not dsRed) signal, therefore the overlap between the reporters negatively correlates with the rate of crinophagic degradation, and mature crinosomes appear as dsRed-positive but GFP-negative structures [30]. We used RNAi-mediated knock-down of *Patsas* and *Hip14* to investigate their putative effect on the crinophagic flux. In contrast to the control cells which contain mostly dsRed-only crinosomes (Fig 1A,D), *Patsas* (Fig 1B) and *Hip14* (Fig 1C) silenced cells accumulate double-positive intact glue granules (Fig 1D). Therefore, the presence of the Patsas and Hip14 PATs is similarly required for the proper crinophagic degradation of glue granules.

**Figure 1.**
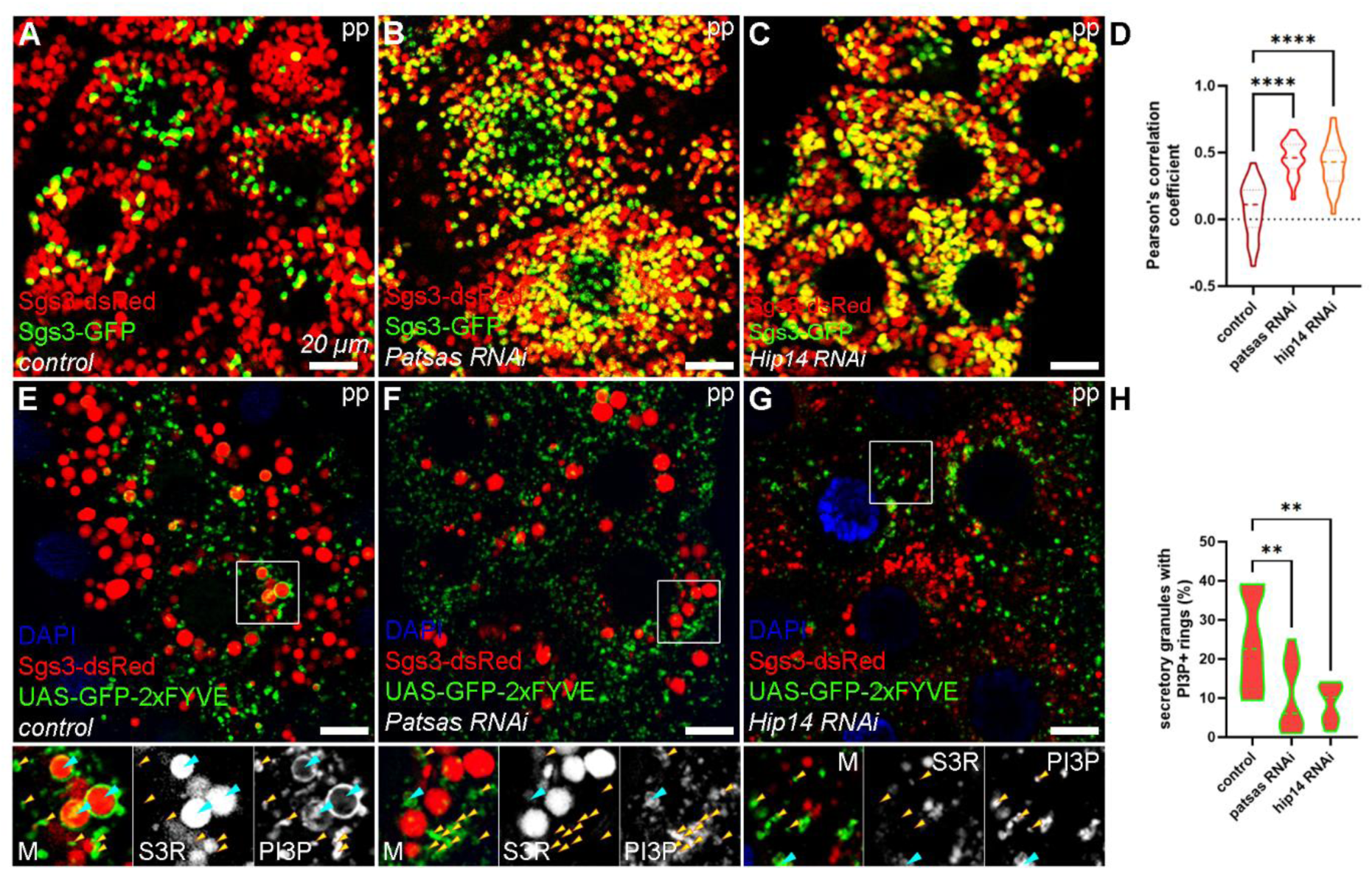
Patsas and Hip14 are essential for proper crinophagic degradation. **(A)** In control prepupal (pp) cells, the Sgs3-GFP signal is quenched in the acidic crinosomes. In contrast, the Sgs3-GFP signal is preserved in salivary glands of same age expressing Patsas **(B)** or Hip14 **(C)** RNAi. **(D)** Quantification of the overlap between the GFP- and dsRed-tagged Sgs3 reporters from **(A-C)**, n=25 cells from 5 different larvae, dashed lines mark the median and dotted lines label the upper and lower quartiles of violin plots, ****p<0.0001. **(E)** In control cells the formation of the crinosomes is supported by fusions between PI3P-positive endosomes (yellow arrowheads) and residual glue granules, resulting in the appearance of GFP-FYVE PI3P-specific reporter in a ring-like pattern around the crinosomes (turquoise arrowheads). In the absence of Patsas **(F)** or Hip14 **(G)** these endosomal fusions are equally decreased and PI3P-positive endosomes accumulate among the glue granules. **(H)** Quantification of the data shown in **(E-G)**, n=353 **(E)**, n=428 **(F)**, n=513 **(G)** glue secretory granules from 3 cells of 3 different larvae, **p<0.01. Insets show 2x magnification of the outlined area, split into channels.

The glue granules at the prepupal stage acquire phosphatidylinositol 3-phosphate (PI3P)-positive membranes presumably through fusion with endosomes, supporting the formation of crinosomes [25]. Along with the Sgs3-dsRed SG marker, we expressed the GFP-myc-2xFYVE PI3P-specific probe to investigate these endosomal fusions. In control cells, FYVE-positive rings appear around the Sgs3-dsRed-positive crinosomes (Fig 1E), indicating successful fusions with PI3P-positive endosomes. However, in the lack of Patsas (Fig 1F) and Hip14 (Fig 1G) PAT enzymes, these endosomal fusions are equally perturbed, and the PI3P+ endosomes accumulate among the glue granules, and the ring-like pattern of FYVE-GFP around the granules mostly disappears (Fig 1H). Thus, Patsas and Hip14 are similarly required for the fusion between the residual glue granules and PI3P-positive endosomes and consequently for the formation of crinosomes.

### 2. Patsas and Hip14 are required for the fusion of maturing secretory granules and lysosomes and their subsequent acidification

Since both Patsas and Hip14 proved to be required for the efficient crinophagic degradation of glue granules, we aimed to test their potential effect on the maturation of the glue granules, preceding their bulk release. The fusion of the secretory granules with a heterogenous lysosomal compartment containing non-degradative lysosomes accompanies the complex maturation process of secretory granules, establishing their proper membrane composition and content for exocytotic release at the age of 2 hours before puparium formation (2h bpf) [24–26]. We analyzed the fusion between maturing glue granules and Arl8-positive lysosomes in salivary gland cells expressing the N-terminally GFP-tagged Arl8 and the Sgs3-dsRed reporters. In control cells, GFP-Arl8 forms rings around the secretory granules indicating successful fusions (Fig 2A). In contrast, in the absence of Patsas (Fig 2B) or Hip14 (Fig 2C), Arl8-positive lysosomes accumulate among the glue granules, rather than forming rings around them (Fig 2D). Therefore, Patsas and Hip14 are similarly required for efficient fusion between maturing secretory granules and lysosomes.

**Figure 2.**
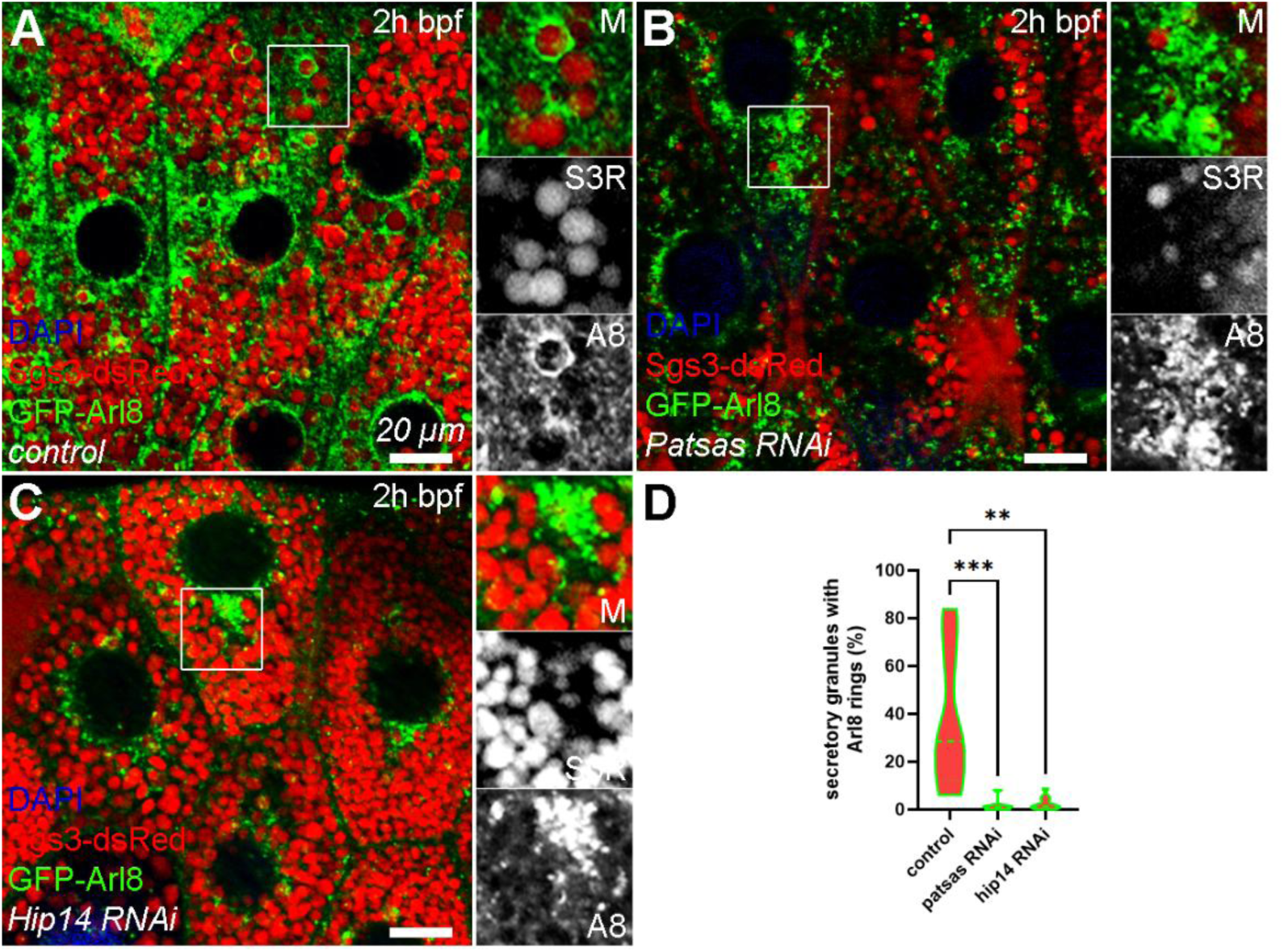
Patsas and Hip14 are required for fusion between Arl8-positive lysosomes and maturing glue secretory granules. **(A)** In control cells Arl8-positive lysosomes fuse with maturing glue granules, resulting in the ring-like appearance of the reporter around their perimeter. In contrast, in the cells lacking either Patsas **(B)** or Hip14 **(C)**, Arl8-positive lysosomes accumulate among the maturing glue granules. **(D)** Quantification of the data shown in **(A-C)**, n=395 **(A)**, n=1162 **(B)**, n=971 **(C)** glue granules from 3 cells of 3 different larvae, dashed lines mark the median and dotted lines label the upper and lower quartiles of violin plots, **p<0.01, ***p<0.001. Insets show 2x magnification of the outlined area, split into channels.

The gradual acidification accompanies the maturation of secretory granules, promoting the remodeling and processing of their secretory content to reach the appropriate consistency before exocytosis [23, 24, 26, 35, 36], and this acidification is highly dependent on secretory granule-lysosome fusions [25]. To assess whether the acidification of maturing glue granules is affected by ANK PATs, we stained salivary gland cells expressing Lamp1-GFP lysosomal reporter with LysoTracker Red vital dye that selectively labels acidic structures. In control cells, large LTR-positive structures of the size of glue granules appear, in which the Lamp1-GFP signal, which usually accumulates in a ring-like pattern around glue granules, is quenched because the GFP moiety located on the luminal side of this reporter is exposed to the acidic milieu (Fig 3A) [37]. However, in the lack of Patsas (Fig 3B) or Hip14 (Fig 3C), the LTR-positive structures are smaller (Fig 3E) and overlap with Lamp1-GFP, representing fusion defective and less acidic lysosomes. As the loss of ANK PATs caused an obvious defect in lysosome fusion and granule acidification, we were also interested in evaluating whether overactivation of Golgi palmitoylation may lead to an opposite effect. To assay this, we overexpressed Hip14 in similarly aged salivary glands that led to a striking expansion of the LTR-positive compartment (Fig3D and 3E). Therefore, we conclude that Patsas and Hip14 are required for proper acidification of lysosomes and thereby maturing secretory granules, and the extent of acidification positively correlates with the level of Hip14.

**Figure 3.**
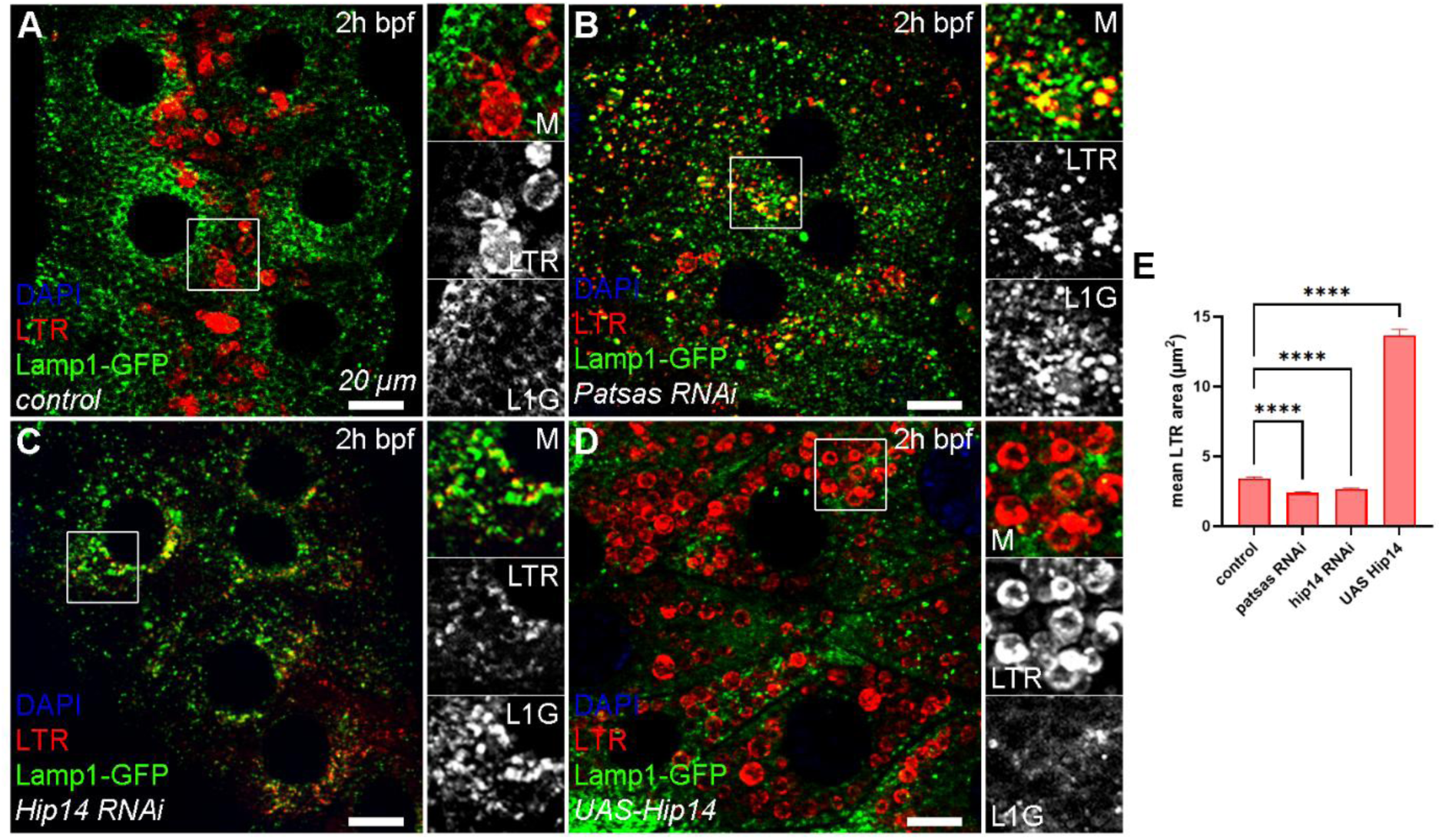
Patsas and Hip14 are required for proper lysosomal acidification. **(A)** In control salivary gland cells large LysoTracker Red-positive (LTR+) acidic structures of the size of glue granules appear, in which the GFP signal of the Lamp1-GFP reporter is quenched. **(B-C)** In the absence of Patsas **(B)** or Hip14 **(C)**, smaller LTR+ structures appear with retained GFP fluorescence. **(D)** In contrast, the overexpression of Hip14 causes the enlargement of LTR+ acidic structures. **(E)** Quantification of the size of the LTR+ structures in **(A-D)**, n=2182 **(A)**, n=2199 **(B)**, n=3853 **(C)**, n=2921 **(D)** LTR+ structure from 5 cells of 5 different larvae, error bars mark ± SEM, ****p<0.0001. Insets show 2x magnification of the outlined area, split into channels.

Acidification, chloride and calcium ion uptake drive the structural remodeling of the content of maturing glue granules and crinosomes, which can be monitored at the ultrastructural level [23, 24, 26]. In control cells 2 hours before puparium formation (2h bpf) large mature granules can be observed with multiple electron-dense cores (Fig 4A). Although in the lack of Patsas (Fig 4B) or Hip14 (Fig 4C) the glue granules show similar morphology, fusion incompetent or docked lysosomes frequently appear in their vicinity. In contrast, salivary gland cells overexpressing Hip14 show small immature secretory granules with the premature presence of large crinosomes with an unusual multivesicular or multilamellar morphology (Fig 4D), likely representing the enlarged acidic structures seen with LTR (Fig 3D). These results further support that Patsas or Hip14 PAT enzymes are required for and act as a rate limiting factors of fusions between maturing secretory granules and lysosomes.

**Figure 4.**
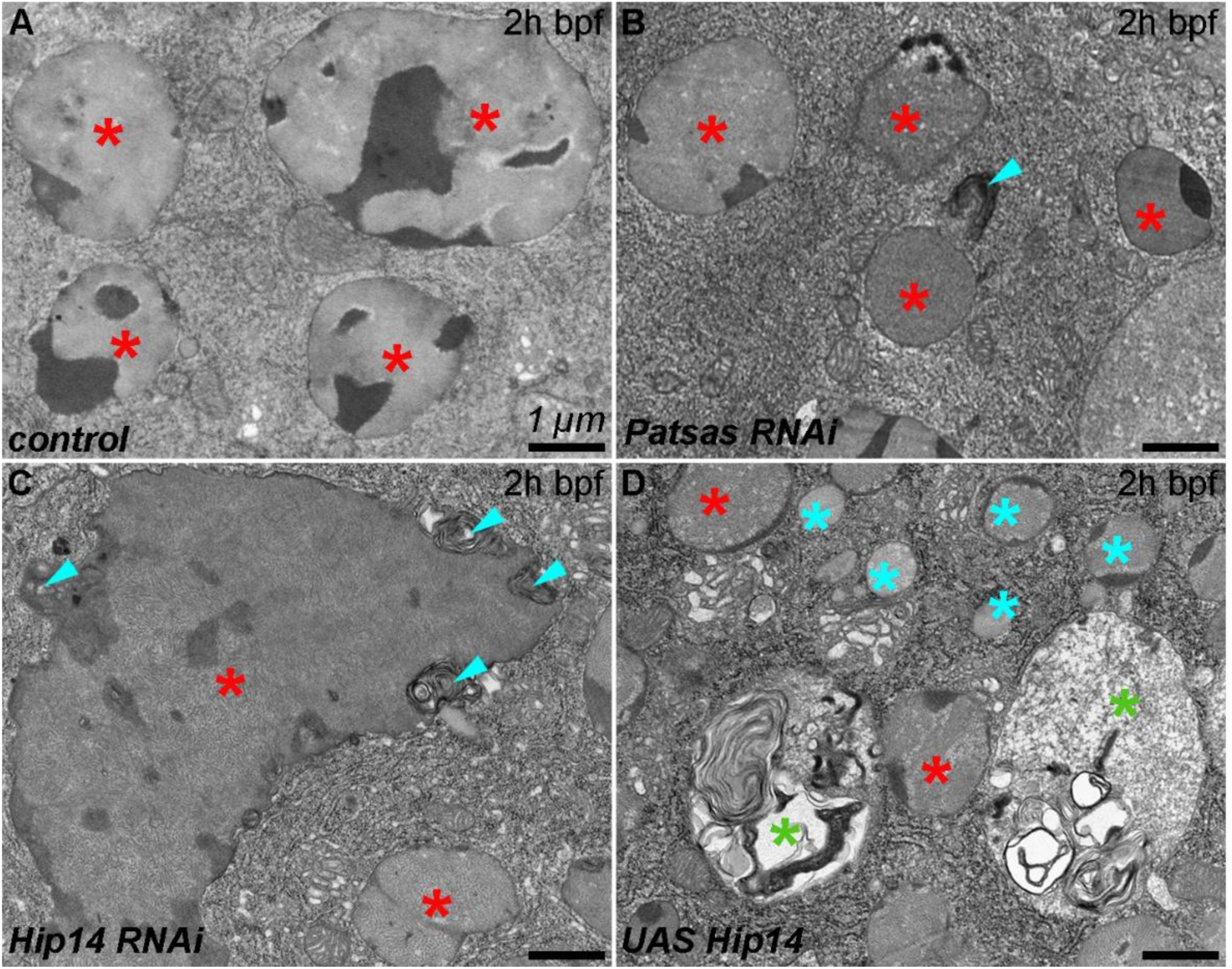
Patsas and Hip14 are required for the effective fusion of lysosomes with maturing glue granules. **(A)** In control cells, large maturing glue granules (red star) with multiple electron-dense cores can be observed shortly before bulk glue secretion (2h bpf). **(B-C)** In same aged salivary glands that are deficient for Patsas **(B)** or Hip14 **(C)**, mature glue granules appear with multilamellar (fusion incompetent) lysosomes (turquoise arrowheads) at their edge. **(D)** In cells overexpressing Hip14 numerous small-sized immature glue granules (turquoise star) appear, together with a premature appearance of abnormal multivesicular crinosomes (green star).

### 3. Patsas and Hip14 are involved in the biosynthetic transport of lysosomal hydrolases

As the presence of both of these Golgi resident ANK PATs was critical for proper fusion of lysosomes with secretory granules, we were interested in whether these enzymes are required for lysosomal biogenesis too. To test the putative role of Patsas and Hip14 enzymes in the sorting and transport of lysosomal proteins, we immunostained the lysosomal protease Cathepsin L in cells expressing the Lamp1-GFP reporter that labels late endosomes and non- or moderately acidic lysosomes [37]. In control cells Cathepsin L showed moderate colocalization with Lamp1-GFP especially on smaller vesicles (Fig 5A). In contrast, the lack of Patsas (Fig 5B) or Hip14 (Fig 5C) equally led to the fragmentation of the Cathepsin L compartment (Fig 5E) and resulted in a more extensive colocalization between the two markers. Finally, the overexpression of Hip14 increased the size of Cathepsin L structures (Fig5 D and 5E), similar to the phenotypes seen with LysoTracker Red (Fig 3). However, these enlarged Cathepsin L positive structures did not colocalize with Lamp1-GFP, which suggests that these structures likely represent the very acidic, multivesicular crinosomes (Fig 4D) where the Lamp1-GFP reporter is degraded rapidly. Taking together, our data suggest that although lysosomal hydrolases can reach the lysosomal compartment even in the absence of Patsas or Hip14, the proper transport and subcellular distribution of lysosomal enzymes is critically influenced by ANK PATs.

**Figure 5.**
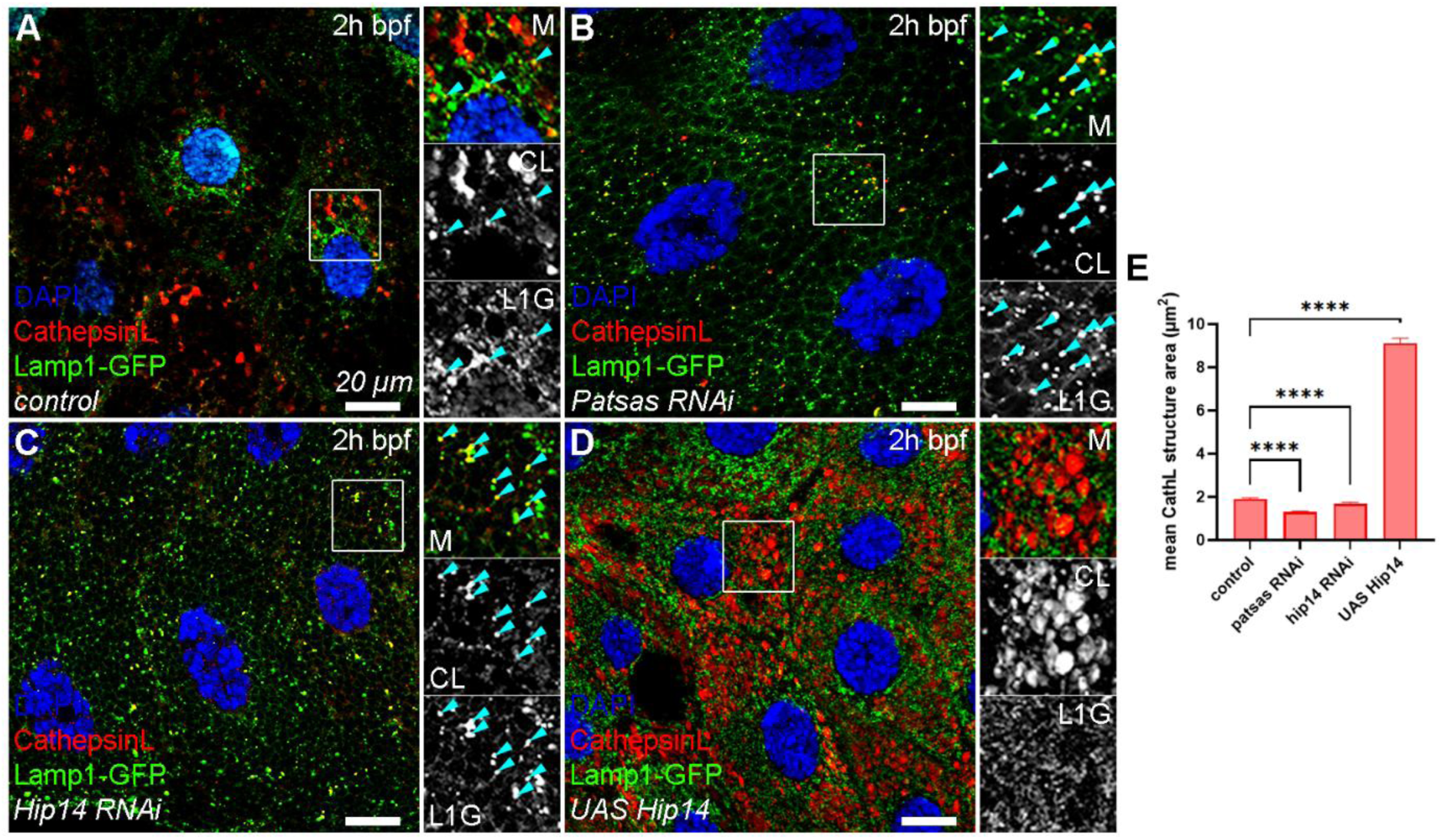
Patsas and Hip14 are essential for proper transport of Cathepsin L lysosomal hydrolase. **(A)** In control cells, the lysosomal protease Cathepsin L colocalize with Lamp1-GFP on small lysosomes. The absence of Patsas **(B)** or Hip14 **(C)** equally led to the fragmentation of the Cathepsin L-positive lysosomal compartment which shows more extensive overlap with Lamp1-GFP. **(D)** In contrast, the overexpression of Hip14 significantly increases the size of Cathepsin L structures, which do not overlap with Lamp1-GFP. **(E)** Quantification of the size of Cathepsin L-positive structures, n=1523 **(A)**, n=788 **(B)**, n=1318 **(C)**, n=1650 **(D)** Cathepsin L+ structures from 5 cells of 5 different larvae, error bars mark ± SEM, ****p<0.0001. Insets show 2x magnification of the outlined area, split into channels.

To follow the transport of lysosomal enzymes at the ultrastructural level, we carried out Gömöri’s acid phosphatase (AcPase) enzyme cytochemistry, that enables us to follow the subcellular localization of lysosomal AcPase enzyme activity. Glue granules gradually acquire the AcPase enzyme during their maturation [24, 26] possibly by fusion with non-degradative lysosomes. We carried out this assay on salivary glands of larvae at different developmental stages. We observed that immature glue granules of control cells at the early L3 larval stage (6h bpf) contain a few acid-phosphatase-positive precipitates (Supplementary Fig 1A). Later, shortly before bulk glue secretion (2h bpf), more AcPase signal is present in mature glue granules (Supplementary Fig 1B). In the lack of Patsas or Hip14 PAT enzymes, the glue granules and the crinosomes contain less AcPase precipitate throughout their maturation (Supplementary Fig 1C-F), accompanied by the simultaneous accumulation of AcPase-containing multilamellar/multivesicular lysosomes at the rim or proximity of the granules at 2h bpf stage (Supplementary Fig 1D and 1F). In contrast, in cells overexpressing Hip14, AcPase-positive crinosomes appear already at the earliest (6h bpf) stage and have an irregular multivesicular morphology (Supplementary Fig 1G). Interestingly, crinosomes that form prematurely fall into two groups: those which mostly contain glue only show more AcPase precipitate, while those with multivesicular morphology, have less but still detectable amount of AcPase precipitate and reduced glue content, presumably due to their enhanced degradation (Supplementary Fig 1G). At the 2h bpf stage, mostly immature glue granules with moderate AcPase signal can be detected and the multivesicular crinosomes have only residual AcPase content (Supplementary Fig 1H). These results demonstrate, that the lysosomal enzyme transport is severely impaired in the lack of Patsas or Hip14 palmitoyl-transferases, which consequently delays the maturation of glue granules too. In contrast, the overexpression of Hip14 was able to enhance the fusion of lysosomes with glue granules, resulting in the premature appearance of Cathepsin L and AcPase-positive crinosomes.

### 4. Expression of the constitutively active form of Rab2 can compensate for the lysosomal defects seen in Patsas or Hip14 deficient cells

The Golgi-localized Rab2 small GTPase protein promotes the biogenesis of the Golgi-derived biosynthetic vesicles and mediates their fusion with lysosomes of different (endocytotic, secretory or autophagic) origin [38–40]. Moreover, overexpressing the constitutively active form of Rab2 was shown to increase glue granule size and their fusion with lysosomes [30]. We sought to examine whether the fragmentation and dysfunctionality of the lysosomal compartment caused by the lack of these PAT enzymes could be restored by ectopic stimulation of lysosomal biogenesis and fusions. Therefore, we expressed a constitutively active Rab2[Q65L] point mutant allele in Sgs3-dsRed expressing salivary gland cells and performed LysoTracker-deepRed (LTdR) staining. In control cells, glue granules are properly acidified and are positive for LTdR (Fig 6A) while, as expected, the expression of the Rab2[Q65L] increased the size of the LTdR-positive glue granules (Fig 6B). Importantly, hyperactivation of Rab2 causes a similarly increased size of these LTdR-positive glue granules even in the absence of Patsas (Fig 6C) or Hip14 (Fig 6D) compared to the control (Fig 6E). Thus, promoting Golgi exit and lysosomal fusions by Rab2 hyperactivation can mediate a compensatory effect on perturbed glue granule acidification caused by ANK PAT deficiency.

**Figure 6.**
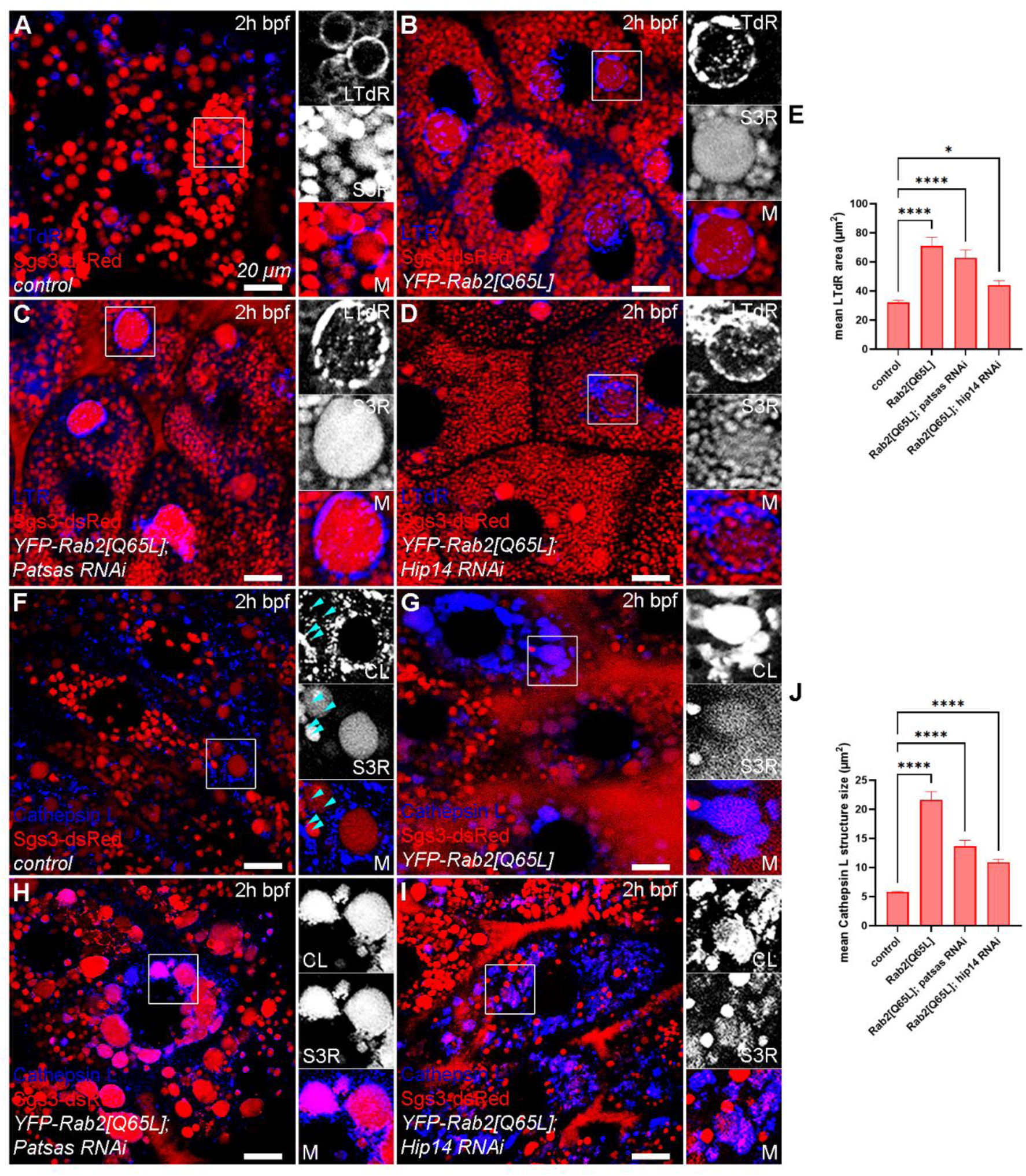
Hyperactivation of Rab2 could compensate for defective acidification and Cathepsin L transport caused by the absence of Patsas and Hip14. **(A)** In control cells maturing glue granules progressively acidify and become positive for LysoTracker DeepRed (LTdR). **(B)** The expression of the constitutively active Rab2[Q65L] transgene increases the size of the LTdR-positive secretory granules. **(C, D)** The expression of Rab2[Q65L] can similarly increase the size of LTdR-positive glue granules even in the absence of Patsas **(C)** or Hip14 **(D)** PAT enzymes. **(E)** Quantification of the size of LTdR-positive structures shown in **(A-D)**, n=150 LTdR-positive structures from 5 cells of 3 different larvae, error bars mark ± SEM, *p<0.05, ****p<0.0001. **(F)** In control cells, Cathepsin L-positive lysosomes appear among the maturing glue granules which also contain some small Cathepsin L-positive foci (turquoise arrowheads). **(G)** The expression of Rab2[Q65L] enhances the fusion of Cathepsin L-positive lysosomes with maturing glue granules and increases their size. **(H, I)** Rab2[Q65L] overexpression can also increase the size of Cathepsin L-positive structures in Patsas **(H)** and Hip14 **(I)** deficient genetic background. **(J)** Quantification of the size of Cathepsin L-positive structures shown in **(G-I)**, n=843 **(F)**, n=765 **(G)**, n=623 **(H)**, n=838 **(I)** Cathepsin L-positive structures from 5 cells of 5 different larvae, **** p<0.0001. Insets show 2x magnification of the outlined area, split into channels.

Next, we investigated whether the fragmentation of Cathepsin L positive compartment in Patsas and Hip14 deficient cells could be similarly reversed by activation of Rab2. Therefore, we carried out immunolabelings with Cathepsin L-specific antibodies on control and Rab2[Q65L] overexpressing salivary glands. In control cells, Cathepsin L-positive lysosomes are mostly positioned among the glue granules, and a few small Cathepsin L positive foci could be also observed inside the granules (Fig 6F) that indicates moderate level of glue granule-lysosome fusion. The expression of Rab2[Q65L] increases the size of the Cathepsin L-containing glue granules by enhancing the fusion of Cathepsin-L positive lysosomes and maturing glue granules, which results in the accumulation of Cathepsin L in these granules (Fig 6G). Importantly, these Cathepsin L-positive granules also increased in size even in the absence of the Patsas (Fig 6H) and Hip14 (Fig 6I) enzymes compared to the control cells (Fig 6F and 6J). These data suggest that the perturbed lysosomal function and impaired glue granule-lysosome fusion in ANK PAT deficient cells (Fig 5B and 5C) could be not only restored, but also overcompensated by ectopically enhancing Rab2 mediated lysosomal biogenesis and fusion.

### 5. Hip14 is required for proper lysosome formation in the adult brain and for adult locomotor performance

As the lack of mammalian Hip14 causes HD-like neurological disorders, we assessed the effect of the absence of Hip14 on the integrity of the lysosomal system in neurons and the neuromuscular function of adult flies. We temporally controlled the expression of *Hip14* RNAi transgene by combining neuron-specific Appl-Gal4 and a temperature-sensitive tub-Gal80 allele [41] to avoid early undesirable lethality [33, 34] and induced the expression of the Hip14 RNAi transgene only in adult flies. First, we immunostained the 14-day-old adult brains with Cathepsin L antibody to analyze the integrity of the lysosomal compartment. Interestingly in brains lacking Hip14, a moderate but significant increase in the size of Cathepsin L-positive structures can be detected (Fig 7A and 7B), indicating that loss of Hip14 leads to dysregulation of lysosome formation in these neurons. We performed a climbing test to examine the effect of neuron-specific loss of Hip14 on adult locomotion. In contrast to the control, the Hip14 knock-down flies show a progressive decline in climbing ability. Importantly, the observed ataxia in Hip14 deficient flies could be restored by the expression of the constitutively active Rab2 transgene (Fig 7C). These results indicate that Hip14 is required for the maintenance of neuronal function, likely through regulating lysosomal formation and function. Furthermore, our results suggest that the ectopic activation of lysosomal biogenesis and fusions could be a good strategy to compensate for the lack of Hip14 and to restore proper neuronal function.

**Figure 7.**
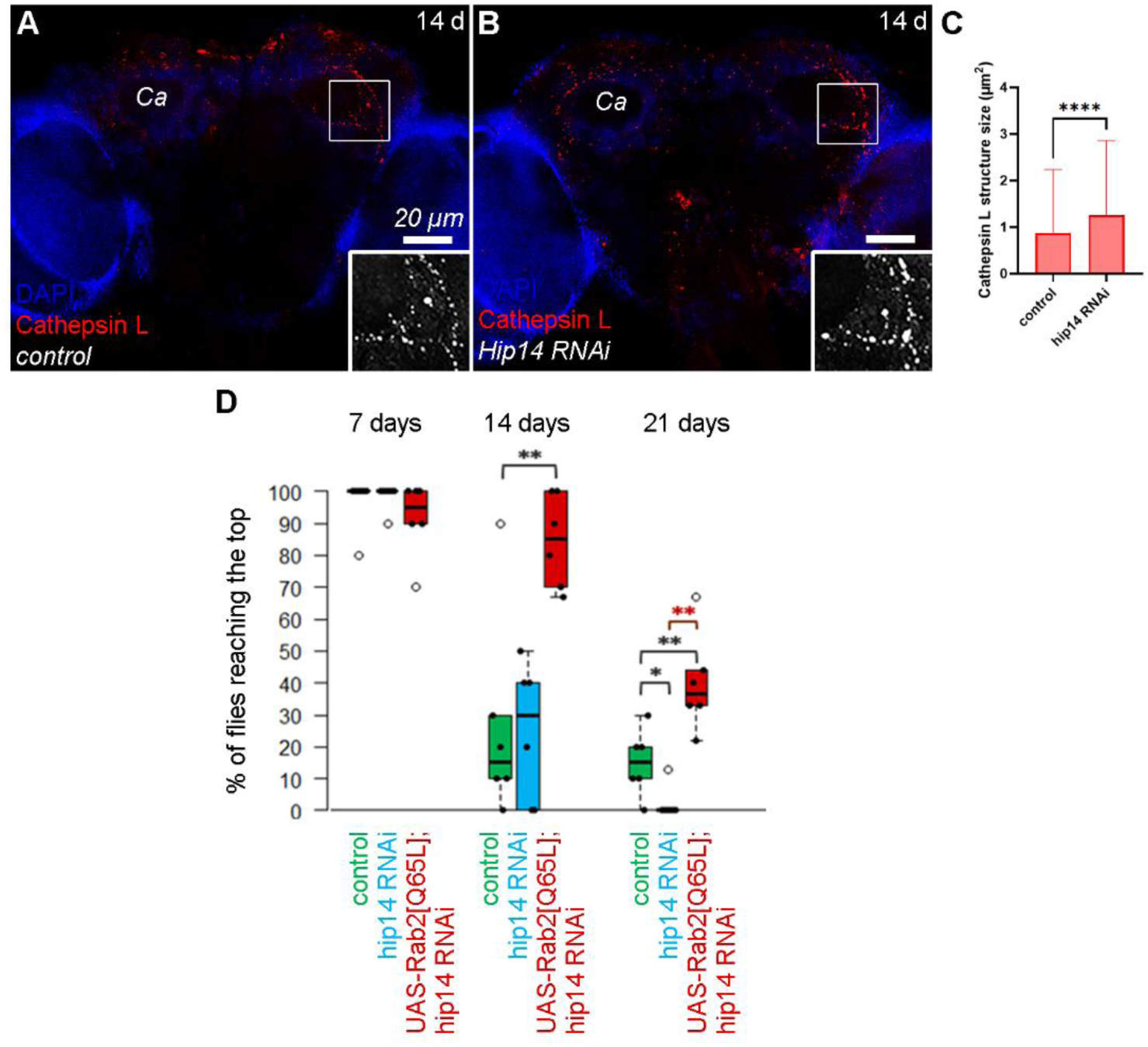
Hip14 is essential for neuronal health and proper lysosomal formation in the adult brains. **(A-B)** Compared to control brains **(A)**, neuron-specific loss of Hip14 leads to a moderate increase in the size of Cathepsin L-positive lysosomes **(B)**. **(C)** Quantification of the size of Cathepsin L-positive lysosomes in 14 days-old adult (female) brains, n=2262 **(A)**, n=1017 **(B)** Cathepsin L-positive lysosomes from 3 different brains, ****p<0.0001. **(D)** The brain-specific knock-down of Hip14 caused a progressive decline in the motor performance of adult flies compared to the control, while the expression of the constitutively active form of Rab2 was able to restore the climbing defect of flies lacking Hip14. Black dots represent one measurement, circles indicate outliers, boxes represent the typical 50% of the climbed adults, the lines show the median and whiskers present the upper and lower quartiles, n=30 flies per each genotype, *p<0.05, **p<0.01. Insets show 2x magnification of the outlined area, split into channels, Ca: calyx neuropil.

## Discussion

In our work, we found that both Patsas and Hip14 are critical regulators of the interplay between lysosomes and the secretory pathway. First, we revealed that both enzymes contribute to efficient crinophagic degradation and fusions between residual glue granules and PI3P-positive endosomes, aiding the formation of crinosomes [25]. Second, we demonstrated that both ANK PATs promote the fusion of maturing glue granules with non-degradative lysosomes, which facilitates their progressive acidification and ultrastructural remodeling [23–26]. Third we found that loss of Hip14 also perturbs lysosomal morphology in the adult brain and leads to gradual impairment in neuromotor performance. Finally, these lysosome-related phenotypes caused by loss of ANK PATs could be partially restored by ectopic activation of lysosomal fusion.

The hypothesis that Patsas and Hip14 can promote lysosomal fusions, and regulate lysosomal protein sorting is further supported by a recent study where Hip14 has emerged as a regulator of *Drosophila* host defense against bacterial infection, presumably by maintaining lysosomal homeostasis [42]. Please note that this recent study did not explore the functionality and fusion capabilities of the affected lysosomes. In our work, we made significant efforts to find out how modulation of ANK PATs affects lysosomal maturation and fusion by assaying the localization of two different lysosomal enzymes. As a result, we demonstrated that loss of either Patsas or Hip14 leads to a significant decrease in fusion between maturing glue granules and Arl8-positive lysosomes and the LysoTracker Red-positive acidic compartment was fragmented, likely representing fusion incompatible lysosomes with a decreased acidification capacity. Moreover, we also observed the fragmentation of the Cathepsin L-positive lysosomal compartment and impaired delivery of lysosomal AcPase to maturing glue granules in ANK PAT deficient secretory cells. Vice versa, the overexpression of Hip14 resulted in the expansion of LysoTracker and Cathepsin L-positive structures, which likely represent premature crinosomes that were also positive for AcPase. These findings suggest that the activity of Golgi resident ANK PATs may serve as a rate-limiting factor in lysosome maturation and fusion. This function of Patsas and Hip14 is further confirmed by our finding that enhancing the fusion of lysosomes with Golgi-derived vesicles or secretory granules in an alternative way [30, 38] – by overactivation of Rab2 GTPase – could compensate for lysosomal dysfunction in Patsas or Hip14 deficient cells. How ANK PATs can regulate lysosomal fusions at the molecular level is still elusive. An important clue might be provided by several studies showing that SNARE proteins [43, 44], and especially the lysosomal R-SNARE Vamp7 [45] – a known critical player in secretory granule-lysosome fusion [25, 30] – can be also palmitoylated. However, how palmitoylation affects Vamp7’s function and whether it is carried out by Patsas and Hip14 is yet to be understood. Additionally, we cannot entirely rule out that the elevated level of lysosomal content in secretory granules in Hip14 overexpressing cells is partially due to the missorting of lysosomal enzymes into the secretory granules, and not only a result of elevated lysosomal fusions. Understanding whether Hip14 has a direct effect on the sorting of lysosomal enzymes in the Golgi requires further research.

We successfully demonstrated that loss of Hip14 also perturbs the lysosomal system in the brain, which is accompanied by declined locomotor performance, a characteristic phenotype of HD-like neurodegeneration [46]. Most importantly, this poor neuromuscular performance could be restored by expressing the constitutively active form of Rab2, consistent with our previous results. A growing amount of evidence suggests that lysosomal dysfunction is a likely cause for a large cohort of neurodegenerative disorders [47, 48], and it was also shown recently that ectopic activation of lysosome fusion in some disease models can ameliorate their symptoms [49]. HD is accompanied by dysfunctional autophagy and defective autophagosome-lysosome fusion [50–52]. Our current research supports that HD-like symptoms can be caused, at least in part, by perturbation of lysosome fusions. Moreover, since HTT was shown to be required for the proper activity of HIP14 and HIP14L [5, 7, 8], our findings that Hip14 and Patsas acts as rate-limiting factors in lysosome fusions may provide a straightforward mechanistic explanation of how mutant HTT leads to lysosomal dysfunction.

One interesting finding of our work is that the loss of Patsas or Hip14 alone is enough to cause a similar perturbation in post-Golgi trafficking and lysosomal fusions. This might be explained by the partially overlapping substrate specificity of the two ANK PATs, and/or the high demand of PAT activity in the Golgi [13], so the cell cannot tolerate the lack of either enzymes. An alternative explanation could be that the activity of Hip14 and Patsas is closely linked to each other, for example one of the enzymes binds substrates stronger but the other performs the palmitoylation more efficiently as suggested for mammalian Golgi PATs [53]. Further research should be conducted to uncover what the actual relationship is between the two ANK PATs’ activities and substrate preference.

In conclusion, our data show that the two Ankyrin-repeat DHHC palmitoyl-transferase enzymes, Patsas and Hip14 are essential for the proper maturation and crinophagic degradation of the secretory granules by mediating their fusion with the endo-lysosomal compartment. Furthermore, the activity of both PAT enzymes is equally required for proper lysosomal function by regulating lysosomal fusion events and mediating the transport of various lysosomal hydrolases. Similarly, the presence of Hip14 is also essential for the proper lysosomal function and neuronal health in the central nervous system, and the defects caused by the absence of Hip14 both in secretory cells and neurons could be partially restored by tissue specific overexpression of constitutively active Rab2. Our results demonstrate that restoration of lysosome biogenesis and fusions may be a good strategy to overcome the lack of the HD-associated PAT, Hip14 and to restore neuronal function.

## Materials and Methods

### Drosophila genetics

The flies were raised at 25 °C temperature on a standard yeast-cornmeal-agar medium. The w^1118^ (#3605), fkh-Gal4 (#78060), appl-Gal4 (#32040), tubP-Gal80^ts^ (#7017), UAS-YFP-Rab2^Q65L^ (#9760), UAS-GFP-myc-2xFYVE (#42712), UAS-Hip14^WT^ (#42697), Sgs3-GFP (#5884) and UAS-patsas^JF01773^ RNAi (#31218) stocks were obtained from Bloomington Drosophila Stock Center (Bloomington, US). The UAS-Hip14^6017R-1^ RNAi line was purchased from NIG-Fly (National Institute of Genetics, Japan). The Sgs3-dsRed reporter was kindly provided by Andrew Andres (University of Nevada, US), the UAS-Lamp1-GFP reporter by Helmut Krämer (University of Texas Southwestern Medical Center, US), UAS-GFP-Arl8 line by Sean Munro (MRC Laboratory of Molecular Biology, UK). To avoid early lethality caused by the absence of *Hip14*, a temperature-sensitive tubP-Gal80 construct was used to regulate the expression of transgenes in the climbing assay temporally. These flies were initially kept at 25 °C temperature during metamorphosis and then the hatched adults were placed at the 29 °C restrictive temperature.

### LysoTracker staining

The larval salivary glands were dissected in cold PBS (pH=7.4) and gently permeabilized for 30 s in 0.05% Triton X-100-PBS (PBTX) solution. The glands were rinsed in PBS (3×30 s) and incubated for 2 min in 0.5 μM LysoTracker Red or 1 μM LysoTracker deep-Red (dissolved in PBS, Invitrogen) solution, then washed in PBS (3×30 s) and mounted with 9:1 PBS: glycerol media containing 1 µg/ml DAPI (4′,6-diamidino-2-phenylindole, Sigma Aldrich) as nuclear stain.

### Immunocytochemistry

The larval salivary glands were dissected in cold PBS, then permeabilized with 0.05% PBTX solution for 30 s and fixed in 4% formaldehyde solution (dissolved in PBS, 40 min, RT). Then, the glands were washed with PBS (3 × 5min, RT), incubated in a blocking solution (5% fetal calf serum in 0.1% PBTX, 30 min, RT) and incubated with the first antibodies (dissolved in blocking solution, overnight, 4 °C). Then the samples were washed (PBTX, 3×10 min) and incubated in blocking solution (30 min, RT) and with the secondary antibodies (diluted in blocking solution, 3h, RT, protected from light). After that samples were incubated in 4% NaCl solution (15 min, RT) which contained Hoechst (1:200, Sigma-Aldrich) nuclear dye and washed (2 × 15 min in 0.1% PBTX, then 3 × 15 min in PBS). Finally, the samples were dissected and mounted in 100% glycerol. The adult brains were fixed in 4% formaldehyde solution (30 min), washed with PBS (3×10 min, RT), blocked (10% fetal calf serum in 0.1% PBTX, 3 hours, RT) and incubated with the first antibodies (in blocking solution) for 2 consecutive nights (4 °C), then with the secondary antibodies (ON, 4 °C). The samples were treated with the Hoechst-containing 4% NaCl solution, washed (same as for salivary glands) and mounted in clear glycerol placed between spacers with the antennal side facing upside.

For the immunostainings chicken anti-GFP (1:1000, Thermo Fisher) and rabbit anti-CathL/MEP (abcam, #ab58991, 1:100) primary and the AlexaFluor488-conjugated anti-chicken and AlexaFluor634- and AlexaFluor568-conjugated anti-rabbit secondary antibodies (Invitrogen, 1:000) were used.

### Fluorescent imaging

Fluorescent images were taken at room temperature with an AxioImager M2 microscope equipped with an ApoTome.2 structured illumination unit (Zeiss, Germany), Orca-Flash 4.0 LT3 digital sCMOS camera (Hamamatsu Photonics, Japan), EC Plan-Neofluar 20x/0.50, Plan-Apochromat 40x/0.95 and Plan-Apochromat 63x/1.4 Oil objectives (Zeiss, Germany). Raw images were processed with ZEN2.3 lite Microscopy Software (Zeiss) and Photoshop CS4 (Adobe Systems, US).

### Electron microscopy

The larval salivary glands and adult eyes were fixed in 1% glutaraldehyde, 2% formaldehyde, 3mM CaCl_2_, 1% D-saccharose in 0.1N sodium cacodylate buffer (pH=7.4, ON, 4 °C). The samples were washed with 0.1N sodium-cacodylate buffer and incubated in 0.5% OsO_4_ (1h, RT) and half-saturated uranyl acetate (30 min, RT), gradually dehydrated in ethanol with increasing concentration and embedded in Durcupan (Fluka) according to the manufacturer’s protocol. The ultrathin (ca. 70nm) sections were contrasted in Reynold’s lead citrate. The samples for Gömöri’s acid phosphatase histochemistry were dissected in 2% glutaraldehyde, 2%formaldehyde containing EM fixative (ON, 4 °C), and the final lead contrasting was omitted, otherwise we followed the protocol described before [54].

### Climbing tests

We measured the climbing ability of the 7, 14 and 21 days old fruit flies (kept at 29°C). Two parallel tubes were used for each measurement. We anesthetized each genotype with CO2, then placed 10-10 females in each tube. The animals then had a rest period of 1.5 hours at 29°C to avoid the aftereffect of CO2. Following a period of rest, the climbing test was repeated three times, with a 30-minute interval between each trial. Each experimental trial was recorded by camera and subsequently analyzed to assess the motor performance of each genotype. The long-term climbing experiment is conducted to the final boundary line (22 cm) on the glass tube, with the objective being to ascertain the number of animals that reach this boundary line within 40 seconds of each genotype [55].

### Statistics

For the quantitative analysis of fluorescent images, the ImageJ software (National Institutes of Health, Bethesda, Maryland, US) was used. The overlap between markers was determined by Pearson’s correlation analysis by using the Coloc2 plugin. The size of the LysoTracker and Cathepsin L structures were analyzed in the 0.5-99 µm^2^ size range to get rid of the background noise and the irrelevant large, coalesced structures with size of multiple glue granules. The threshold for these images were set by the same person. The enlarged Cathepsin L-positive structures were counted in the relevant 3-500 μm^2^ range, while the LysoTracker deep-Red structures were manually selected. For the quantitative assessment of ring-like structure formation, the glue granules were manually selected and the percentage of them with ring-like structures was analyzed. The small lysosomes in adult brains were analyzed in 0.1-10 µm^2^ range. For comparison of datasets following Gaussian distribution, one-way ANOVA with Tukey’s post-hoc test (Fig 1D and 1H) and if at least one of the datasets followed a non-Gaussian distribution, Kruskal-Wallis test (Fig 2D, Fig 3E, Fig 5E, Fig 6E and 6J) was performed. For pairwise comparison of datasets following non-Gaussian distribution, Mann-Whitney U-test was performed (Fig 7C, Fig 8D). The statistical analyses were performed with the GraphPadPrism 9.0.0 software (Boston, Massachusetts, US) and for the climbing tests with RStudio (v2023.06.2).

## Acknowledgements

We thank Ivett Répássy and Sarolta Pálfia for technical assistance, and colleges and stock centers for providing reagents.

## Supplementary Figures and Supplementary Figure Legends

**Supplementary Figure 1.**
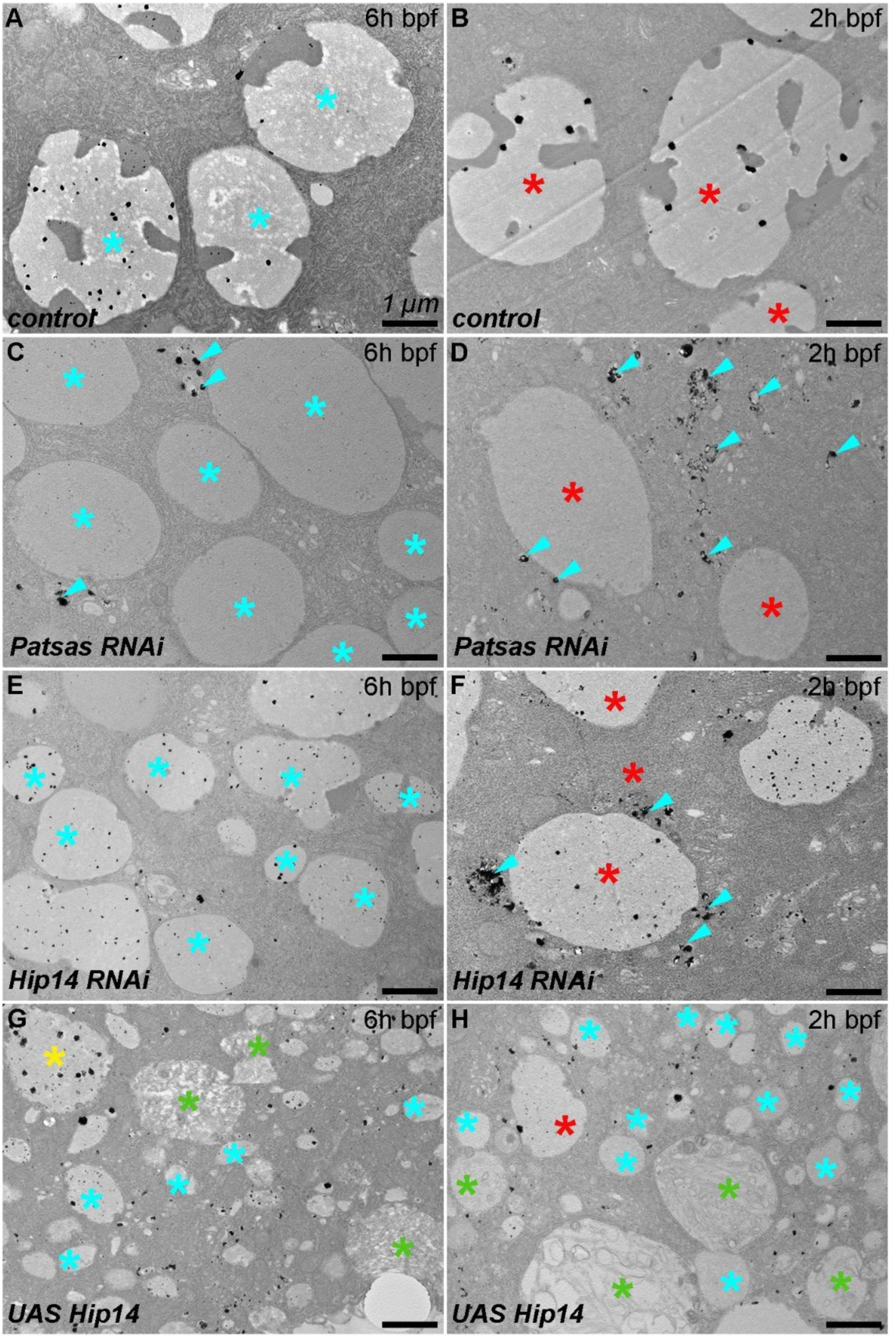
Patsas and Hip14 are required for proper transport of acidic phosphatase enzyme into maturing glue secretory granules. **(A)** The small immature granules (turquoise star) in control cells of wandering (6h bpf) larvae contain very few acidic phosphatase (AcPase) precipitate (appears as black dots). **(B)** The mature glue granules (red star) shortly prior to the bulk glue secretion (2h bpf) contain more AcPase. **(C)** In the absence of Patsas, immature glue granules contain similar level of AcPase signal to the age-matched control, while highly AcPase-reactive lysosomes (turquoise arrowheads) can be observed in the proximity of the granules. **(D)** 2h bpf stage mature glue granules have a moderate AcPase signal compared to the control, while AcPase-containing (fusion incompetent) lysosomes accumulate among them. **(E)** The immature glue granules of cells lacking Hip14 show similar AcPase enzyme activity to the control, while the mature glue granules at 2h bpf stage contain slightly less which is accompanied by the appearance of highly AcPase-positive lysosomes at their close proximity **(F)**, similar to the *Patsas* mutant **(D)**. **(G)** In contrast, the immature glue granules of Hip14 overexpressing cells contain a lot of AcPase precipitate, with the premature appearance of crinosomes. These crinosomes split into two groups, those with the conventional content containing much (yellow star) and those with an irregular multivesicular morphology (green star) with less AcPase positivity. **(H)** Later at the 2h bpf stage Hip14 overexpressing cells contain AcPase-positive immature glue granules.

